# Bromodomain 4 inhibition leads to *MYCN* downregulation in Wilms’ tumor cells

**DOI:** 10.1101/2021.01.04.425208

**Authors:** Andrew D. Woods, Noah E. Berlow, Reshma Purohit, Katherine E. Tranbarger Freier, Joel E. Michalek, Melvin Lathara, Kevin Matlock, Ganapati Srivivasa, Brigitte Royer-Pokora, Renata Veselska, Charles Keller

**Affiliations:** Children’s Cancer Therapy Development Institute, Beaverton, OR 97005 USA; Department of Population Health Sciences, Joe R. & Teresa Lozano Long School of Medicine, University of Texas Health Science Center, San Antonio, TX 78229 USA; Omics Data Automation Inc., Beaverton, OR, USA; Institute of Human Genetics, Medical Faculty, Heinrich-Heine-University Duesseldorf, Germany; Department of Experimental Biology, Faculty of Science, Masaryk University, 61137 Brno, Czech Republic

**Author notes:** correspondence: Charles Keller MD, 12655 SW Beaverdam Rd W, Beaverton OR 97005 USA.

**Keywords:** Anaplasia, AZD5153, *BRD4*, *MYCN*, Wilms’ tumor

## Abstract

Wilms’ tumor is the most common childhood kidney cancer. Two distinct histological subtypes of Wilms’ tumor have been described: tumors lacking anaplasia (the favorable subtype) and tumors displaying anaplastic features (the unfavorable subtype). Children with favorable disease generally have a very good prognosis, while those with anaplasia are oftentimes refractory to standard treatments and suffer poor outcomes. *MYCN* dysregulation has been associated with a number of pediatric cancers including the anaplastic subtype of Wilms’ tumor. In this context, we undertook a functional genomics approach to uncover novel therapeutic strategies for those patients with anaplastic Wilms’ tumor. Genomic analysis and *in vitro* experimentation demonstrate that Wilms’ tumor cell growth can be reduced by modulating *MYCN* overexpression via BRD4 inhibition. We observed a time dependent reduction of MYCN and MYC protein levels upon BRD4 inhibition in Wilms’ tumor cell lines which led to increased cell death and suppressed proliferation. We suggest that AZD5153, a novel dual-BRD4 inhibitor, can reduce MYCN levels and should be further explored for its therapeutic potential against Wilms’ tumor.

## Introduction

Nephroblastoma (Wilms’ tumor) is the most common pediatric kidney cancer, affecting approximately 500 and 1000 children annually in North America and Europe, respectively (Howlader N, Gatta et al., 2014), and overall 1 in 10,000 children worldwide (Breslow et al., 1993). The disease was first described in detail by German pathologist/surgeon Carl Max Wilhelm Wilms in his 1899 seminal paper, *Die Mischgeschwülste der Niere* (The Mixed Tumors of the Kidney) (Wilms, 1899). Wilms’ tumor is thought to arise from the malignant transformation of abnormally persistent prenatal renal stem cells that retain embryonic differentiation potential (Breslow and Beckwith, 1982). Management of the disease has been hailed as a model of modern-day multimodal treatment success, achieving overall cure rates greater than 90% (Dome et al., 2015). The vast majority (90%) of Wilms’ tumor patients are described as having favorable histology (lacking anaplasia). Patients with favorable Wilms’ tumor tend to respond positively to standard therapeutic approaches such as surgery, chemotherapy and radiation. However, approximately 10% of Wilms’ tumor patients have tumors that display varying degrees of anaplastic features such as large nuclei, atypical miotic features, and abnormal patterning due to a loss of polarity. These patients make up the unfavorable subtype of the disease and tend to experience resistance to treatment and suffer poor outcomes (Phelps et al., 2018, Dome et al., 2006, Ooms et al., 2016). The 10-year overall survival rate (OS) for children with stage 4, anaplastic Wilms’ tumor is 18%, versus an 81% 10-year OS for children with stage 4 non-anaplastic Wilms’ tumor (Davidoff, 2012). Therapeutic strategies for children with unfavorable Wilms’ tumor include multiagent high-dose chemotherapy and radiation, leaving survivors with a variety of life-long sequelae. Novel therapies tailored specifically to the unfavorable subtype of Wilms’ tumor are therefore warranted to address the present day unmet clinical need.

While outcomes and response to treatments differ greatly between the two subtypes of Wilms’ tumor, very few significant differences between the favorable and unfavorable groups have been identified at the genomic level. The most common observation has been *TP53* mutations associated with the anaplastic subtype (Bardeesy et al., 1994). *TP53* mutations are often observed within anaplastic regions of the heterogeneous tumor, but rarely observed within non-anaplastic regions (Ooms et al., 2016). In 2005, Li et al. performed transcript profiling on 54 Wilms’ tumors (8 unfavorable) and identified 28 genes differentially expressed between the anaplastic and non-anaplastic groups (Li et al., 2005). However, many of these genes are involved with nuclear integrity, apoptosis and proliferation, suggesting that differences in expression profiles might merely be reflective of the aggressive nature of anaplastic disease. In 2015, Williams et al. observed an association between the anaplastic subtype of Wilms’ tumor and *MYCN* copy number gain which was associated with decreased overall survival (Williams et al., 2015). Thirty percent of diffuse anaplastic Wilms’ tumor samples had *MYCN* copy number gain in the Williams study. *MYCN* gain or *MYC* amplification has been observed in several pediatric cancers, including neuroblastoma (Pugh et al., 2013, Campbell et al., 2017), rhabdomyosarcoma (Williamson et al., 2005), and medulloblastoma (Pfister et al., 2009). In fact, *MYC* dysregulation has been observed in >50% of all human cancers.

The *MYC* family of cellular oncogenes includes family members *MYCC, MYCN*, and *MYCL. MYC* is a pleiotropic basic helix–loop–helix leucine zipper transcription factor involved in many oncogenic processes, including proliferation, apoptosis, differentiation, cell adhesion, signal transduction, translation/transcription, DNA repair and metabolism (Chen et al., 2018, Dang et al., 2006, Arvanitis and Felsher, 2006). Direct targeting of *MYC* oncogenes has thus far proven challenging due to a protein structure that lacks specific binding sites for small molecules and *MYC*’s nuclear location (Chen et al., 2018). This has led to indirect targeting strategies, such as *MYC/MAX* complex disruption (Berg et al., 2002), *MYC* transcription and/or translation inhibition (Delmore et al., 2011), and *MYC* destabilization (Tavana et al., 2016). *MYCN* is a member of the *MYC* family of oncogenes and has been associated with a variety of aggressive cancer types (Rickman et al., 2018). *MYCN* has been implicated in metastasis, survival, proliferation, pluripotency, self-renewal, and angiogenic pathways (Huang and Weiss, 2013) and has been linked to the development of chemoresistance in neuroblastoma (Carter et al., 2015, Huang and Weiss, 2013). *MYCN* has also been shown to play a role in normal kidney nephron development (Bates et al., 2000). In the present study, we focus on *MYC* and*MYCN*dysregulation in Wilms’ tumor, and suggest that the *MYCN* overexpression observed in anaplastic Wilms’ tumor can be downregulated by targeting *MYC* transcription via bromodomain 4 (BRD4) inhibition.

*BRD4* is a transcription co-activator and member of the mammalian bromodomain and extraterminal (BET) family of chromatin readers, characterized by tandem bromodomains (BD1 and BD2) which bind acetylated histones and transcription factors (Yang et al., 2005). BRD4 is known to regulate *MYC* transcription by controlling chromatin-dependent signal transduction to RNA polymerase (Nucera, 2019), increasing the effective concentration of recruited transcriptional activators (Delmore et al., 2011) and directly interacting with the Positive Transcription Elongation Factor Complex b (P-TEFb) (Yang et al., 2005). *BRD4* is known to predominantly bind enhancer elements important for cell type specification and oncogenesis (Lovén et al., 2013). Pharmacological inhibition of BRD4 has demonstrated *in vitro* and *in vivo* tumoristatic effects in a variety of cancer types, including acute myeloid leukemia (Zuber et al., 2011), diffuse large B cell lymphoma (Chapuy et al., 2013), prostate cancer (Asangani et al., 2014) and breast cancer (Shu et al., 2016). In the present study, we show that BET/BRD inhibition via a BRD4 inhibitor results in lowered levels of both MYC and MYCN proteins in Wilms’ tumor cell lines, leading to *in vitro* cell death and proliferation suppression.

## Results and Discussion

### BET/BRD inhibitors attenuate growth of anaplastic Wilms’ tumor cell lines

In the discovery phase of our study, we conducted exploratory chemical screens against a variety of anaplastic and non-anaplastic Wilms’ tumor cell lines and primary cultures (**Fig 1A, Fig EV1**). In our initial 60-compound screen, we identified several inhibitors that showed cell killing activity against all Wilms’ tumor models. We chose to focus on BRD4 inhibition because both BRD4 inhibitors within our screen (JQ1 and I-BET-762) showed a trend of activity against the anaplastic models and only minimal activity against the non-anaplastic models (**Fig 1A**). Based on our initial observation that BRD4 inhibition was effective at reducing anaplastic Wilms’ tumor cell viability, we next wanted to determine the most effective BET/BRD inhibitor against our Wilms’ tumor cultures. We tested 5 different BET/BRD inhibitors against 3 Wilms’ tumor cell lines (**Fig 1B**). Within these screens, AZD5153 was the most efficacious drug against all 3 cell lines. Next, AZD5153 was evaluated against an expanded panel of Wilms’ tumor cell cultures (**Fig 1C**) and IC50 values ranging from 140nM to 2.1μM were observed. Importantly, AZD5153 was 4-fold less potent against normal kidney cell line HEK293 (IC50=4.37μM) than Wilms’ tumor samples (median IC50=1.1μM).

**Figure 1.**
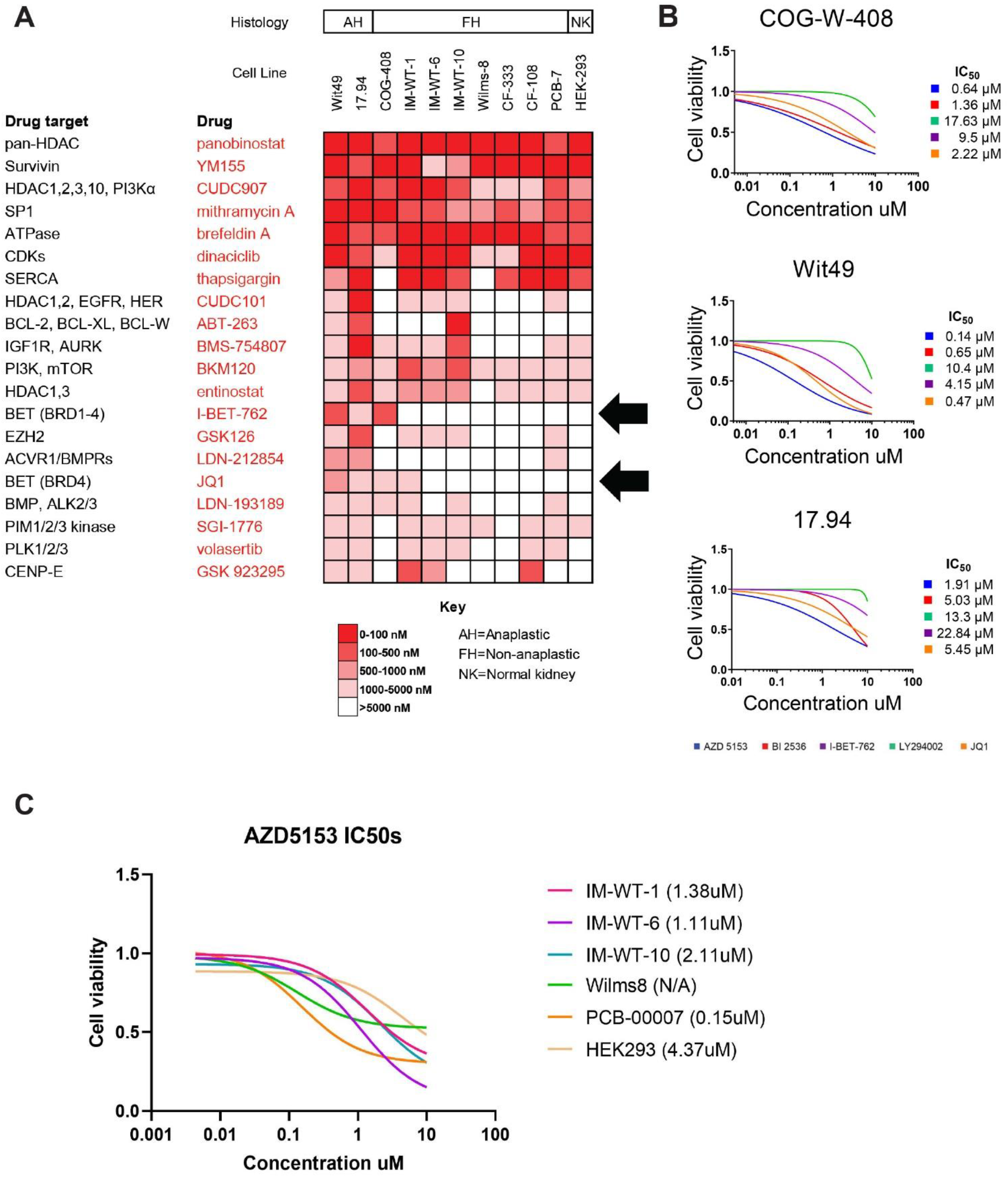
Wilms’ tumor chemical screens. A. Heatmap showing IC_50_ values of 20 inhibitors of a variety of oncogenic pathways. Red indicates strong activity (IC_50_ = <100nM), white indicates no activity (IC_50_ = >5000nM) and scaling between. Arrows highlight BET/BRD inhibitors JQ1 and I-BET-762 which demonstrate preferential activity against anaplastic Wilms’ tumor models. B. Drug curves of 5 BET/BRD inhibitors against 3 Wilms’ tumor cell lines. IC_50_ values were calculated for each of inhibitor. AZD5153 consistently demonstrated the strongest activity (lowest IC_50_ values) against Wilms’ tumor cell lines. C. AZD5153 IC_50_ values across a panel of Wilms’ tumor cell lines and primary cultures. Note the range of effect across samples, with the least effect against HEK293, a normal kidney cell line. Median IC_50_ of Wilms’ tumor samples is 4-fold lower than HEK293 IC_50_ value.

AZD5153 is a potent, selective, orally available, reversible bivalent inhibitor of BRD4 which is currently in phase I clinical trials as a monotherapy and combination therapy for lymphoma, ovarian cancer, and other refractory tumors (NCT03205176 and NCT03527147). Unlike previously developed monovalent BET/BRD inhibitors such as JQ1, AZD5153 distinctly binds and ligates both bromodomains of BRD4 (BD1 and BD2) simultaneously, resulting in increased affinity and potency (Rhyasen et al., 2016). Preliminary data from the AZD5153 first-in-human clinical trial (NCT03205176) indicates that the drug is safe and tolerated at oral doses up to 30 mg QD and 15 mg BID, and displays a dose-dependent linear increase in pharmacokinetics (Wang et al., 2019).

Given that BET/BRD inhibition has been used as a strategy to indirectly regulate *MYC/MYCN* expression in several cancer types, we hypothesized that JQ1’s selective activity within our discovery drug screen was due to *MYCN’s* overexpression in anaplastic Wilms’ tumor models compared to either normal kidney or non-anaplastic models (**Fig 2A**). To evaluate the potential correlation between *MYCN* overexpression and decreased overall survival in Wilms’ tumor, we performed a Cox regression (survival) analysis of data collected from 125 high-risk Wilms tumor patients within the National Cancer Institute’s Therapeutically Applicable Research to Generate Effective Treatments (TARGET) study. We found that increased *MYCN* expression was strongly associated with a decrease in overall survival (P= 0.02) (**Fig 2B**). Interestingly, *MYCN* expression levels had little effect on overall survival during the first 2 years post diagnosis, however, 4 years post diagnosis approximately 75% of the children in the *low-MYCN* group remained alive compared to less than half in the high-*MYCN* group. We did not observe a significant association between high *MYC* expression and overall survival (P= 0.3). Next, we performed a cluster analysis utilizing gene expression data from our Wilms’ tumor samples and a panel of *MYC* associated genes (**Fig 2C**, **Fig EV 2**). Genes chosen for the analysis included *MYC* family genes, *BRD* family genes, *MYC* target genes, *MYC* regulatory genes and genes associated with the *MYC* pathway. Clustering was observed between *MYCN* and *LEF1* genes (**Fig 2D**), indicating that these genes were expressed in similar patterns across our anaplastic and non-anaplastic Wilms’ tumor samples. An ordinary one-way ANOVA analysis showed that *MYCN* and *LEF1* overexpression were strongly associated (P = <0.0001) with the anaplastic subtype of Wilms’ tumor (**Fig 2E**). *LEF1* is known to be regulated by MYC (Hao et al., 2019) and is a transcription co-activator within the Wnt/beta-catenin pathway, a pathway which can promote proliferation and is dysregulated in many Wilms’ tumors (Perotti et al., 2013, Koesters et al., 1999, Maiti et al., 2000, Royer-Pokora et al., 2008). To our knowledge, this observation is the first report of *LEF1* overexpression being linked to the anaplastic subtype of Wilms’ tumor.

**Figure 2.**
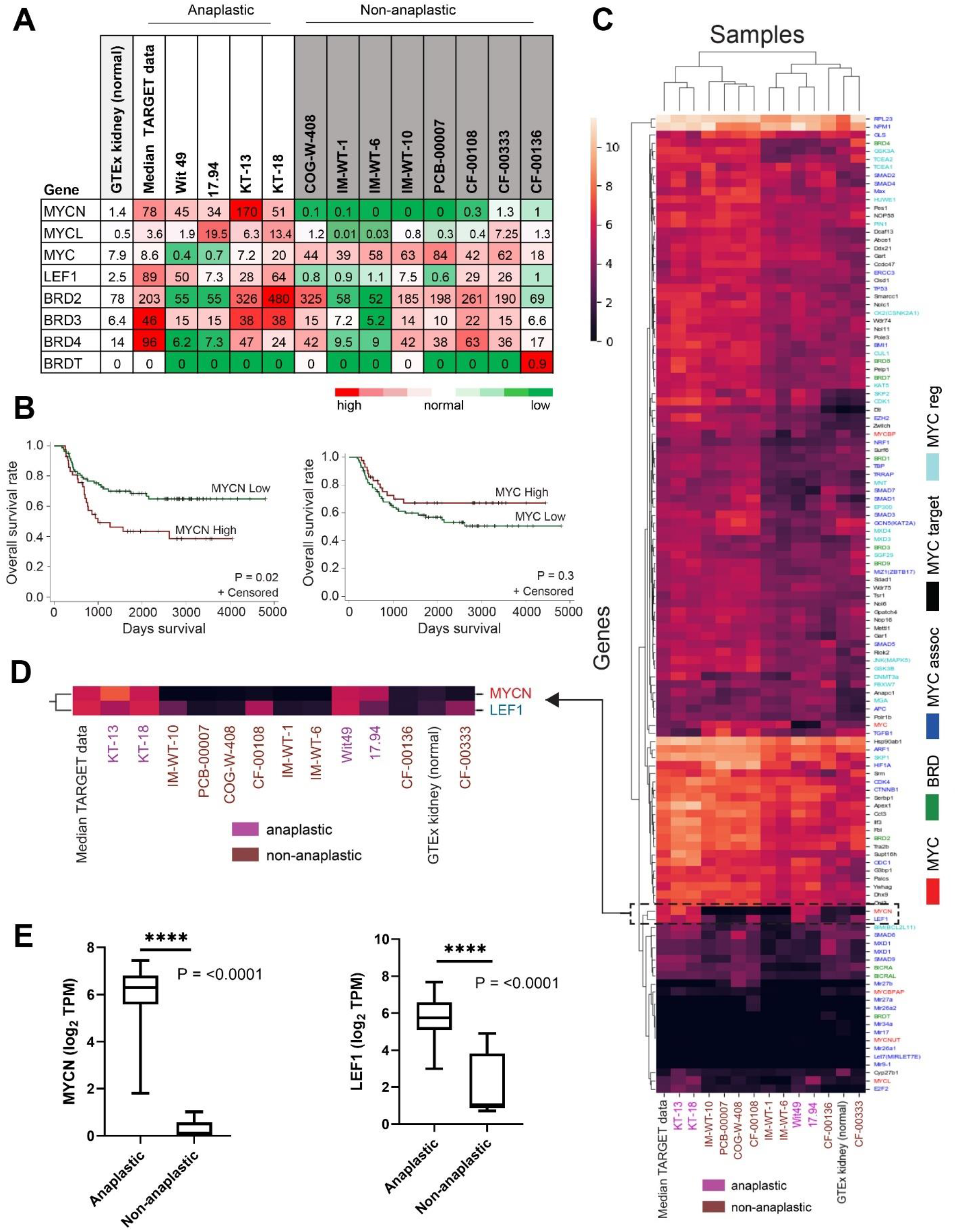
*MYC* and *BRD* gene expression (TPM) in Wilms’ tumor. A. Heatmap showing expression levels of *MYC* and *BRD* genes in a panel of Wilms’ tumor samples as compared to normal kidney. Red indicates overexpression, green indicates underexpression. *MYC* gene family members *MYCN* and *MYCL* and *MYC*-associated gene *LEF1* show an overexpression pattern which is associated with anaplastic Wilms’ tumor. B. Cox regression for survival analysis showing overall survival was significantly shortened in patients with high *MYCN* expression (top third) compared to patients with medium and low *MYCN* expression (P = 0.02). Red is high expression, green is low expression. High *MYC* expression showed no association (P= 0.3) with increased overall survival. Survival data was collected from the NCI TARGET study. C. Cluster analysis and dendrogram heatmap of Wilms’ tumor gene expression levels (TPM) across an expanded panel of *MYC* related genes. D. Breakout of Wilms’ tumor cluster analysis heatmap showing that *MYCN* and *LEF1* genes are expressed in similar patterns between anaplastic and non-anaplastic Wilms’ tumors. E. Box-and-whisker plot representation showing that MYCN and LEF1 are significantly overexpressed (P= <0.0001) in anaplastic Wilms’ tumors (n=42) versus non-anaplastic Wilms’ tumors (n=8). Datasets were analyzed by ordinary one-way ANOVA.

### BRD4 inhibition lowers MYC and MYCN levels in Wilms’ tumor cell lines and increases cell death when combined with radiation

Immunoblotting studies were conducted to determine endogenous MYC protein levels within our Wilms’ tumor samples (**Fig 3A**). To examine the effect of BET/BRD inhibition on MYC protein levels over time, we tested model-specific IC50 concentrations of AZD5153 against three MYC-expressing Wilms’ tumor cell lines. Consistently, a pattern of MYCN and MYC protein reduction at 24 hours of AZD5153 exposure was observed, followed by a recovery of protein levels after 72 hours (**Fig 3B**). To rule out a possible cell cycle effect on *MYCN* levels, untreated cells were collected over the same time course and MYCN protein levels were measured. Consistent MYCN protein levels were observed over the 72-hour time period (**Fig 3C**). To rule out the possibility of decreased drug activity over time, additional experiments were performed in which drug and culture media were refreshed every 24 hours. Again, we saw a reduction of MYC protein levels at 24 hours followed by a rebound of MYC protein levels at 72 hours. One strategy for future study might be to explore ways to extend this period of MYC downregulation. Although MYCN downregulation was not sustained over time in our experiments, other work has noted that even transient inactivation of *MYC* can result in sustained tumor regression in certain cancer types *in vivo* (Arvanitis and Felsher, 2006, Jain et al., 2002).

**Figure 3.**
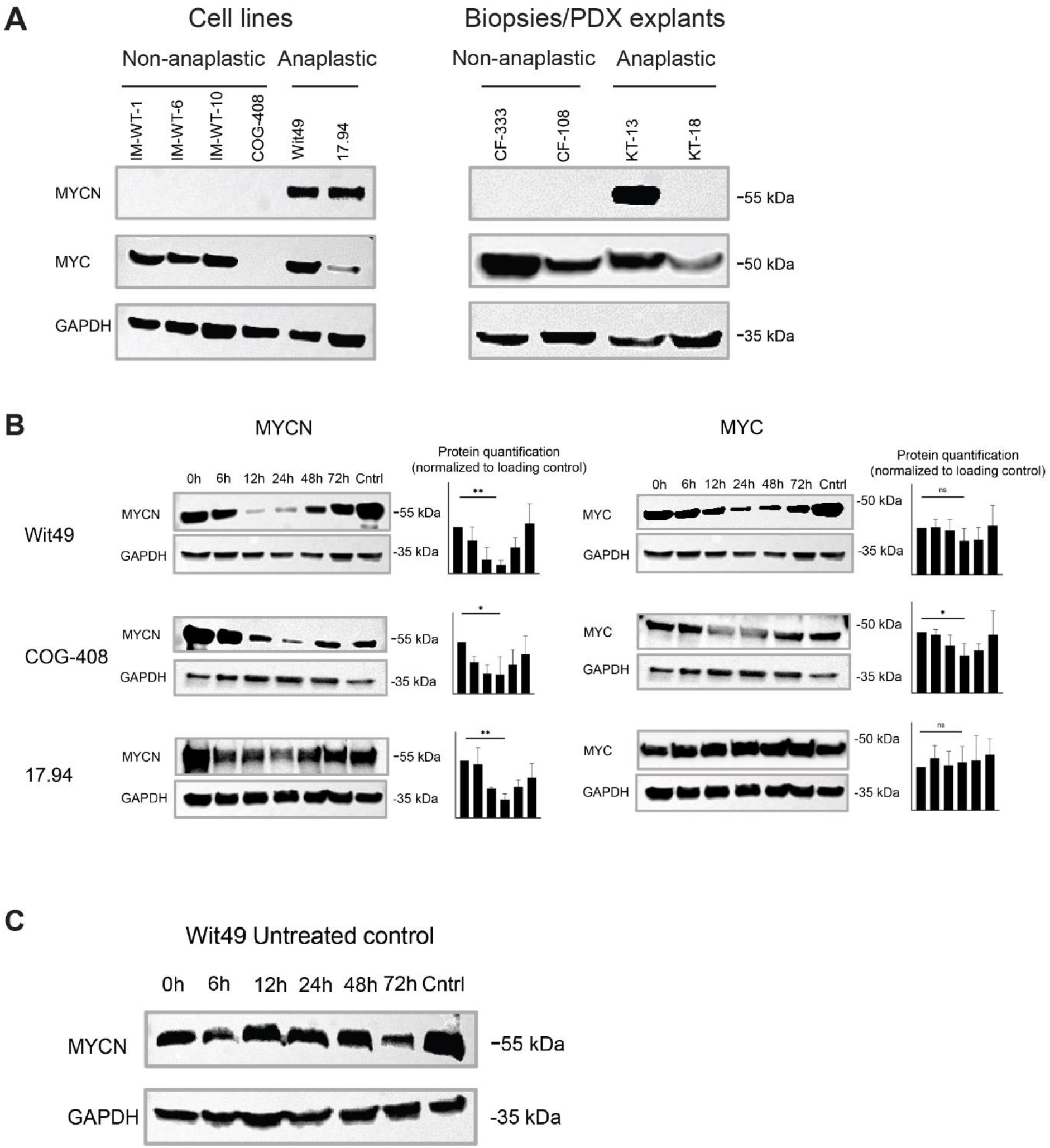
Western blots of MYCN and MYC in Wilms’ tumor. A. Endogenous protein levels of MYCN and MYC in Wilms’ tumor cell lines, biopsies and PDX tissue explants. B. MYCN and MYC levels in Wilms’ tumor cell lines over time in response to BRD4 inhibition by model-specific IC_50_ concentrations of AZD5153. Western blots were performed in triplicate. Each replicate was quantified by number of pixels and then normalized to loading controls. Significance for each cell line was determined by one-tailed t-test of the hypothesis that protein level at 24 hours was less than that at time 0 hours. To account for multiple testing, the Bonferroni approach was used, leading to a test-level alpha of 0.0083 to control the overall alpha to no more than 0.05. All three cell lines met this significance threshold when probing for MYCN. COG-408 met this significance threshold when probing for MYC. C. With no drug treatment, MYCN levels remain constant in Wilms’ tumor cell line Wit49 over a 72hour period.

In order to determine the mode of cell death upon AZD5153 exposure, we performed Annexin V and Propidium Iodide staining experiments with 3 Wilms’ tumor cell lines over a 72-hour period which showed no significant increase in necrotic (dead) and apoptotic (dying) cells in response to AZD5153 treatment over time (**Fig 4A and B**). We then looked at BrdU incorporation as a measure of cellular proliferation and saw a significant reduction of BrdU incorporation (P = <0.0001) upon model-specific IC50 AZD5153 treatment over a 72-hour period (**Fig 4C**). Reduction of BrdU was greater than reduction of cell viability in response to AZD5153 treatment (**Fig 4D**). This suggests that in Wilms’ tumor cells, a major mode of action of AZD5153 is the suppression of cellular proliferation and that BRD4 inhibition by AZD5153 does not necessarily lead to apoptotic cell death.

**Figure 4.**
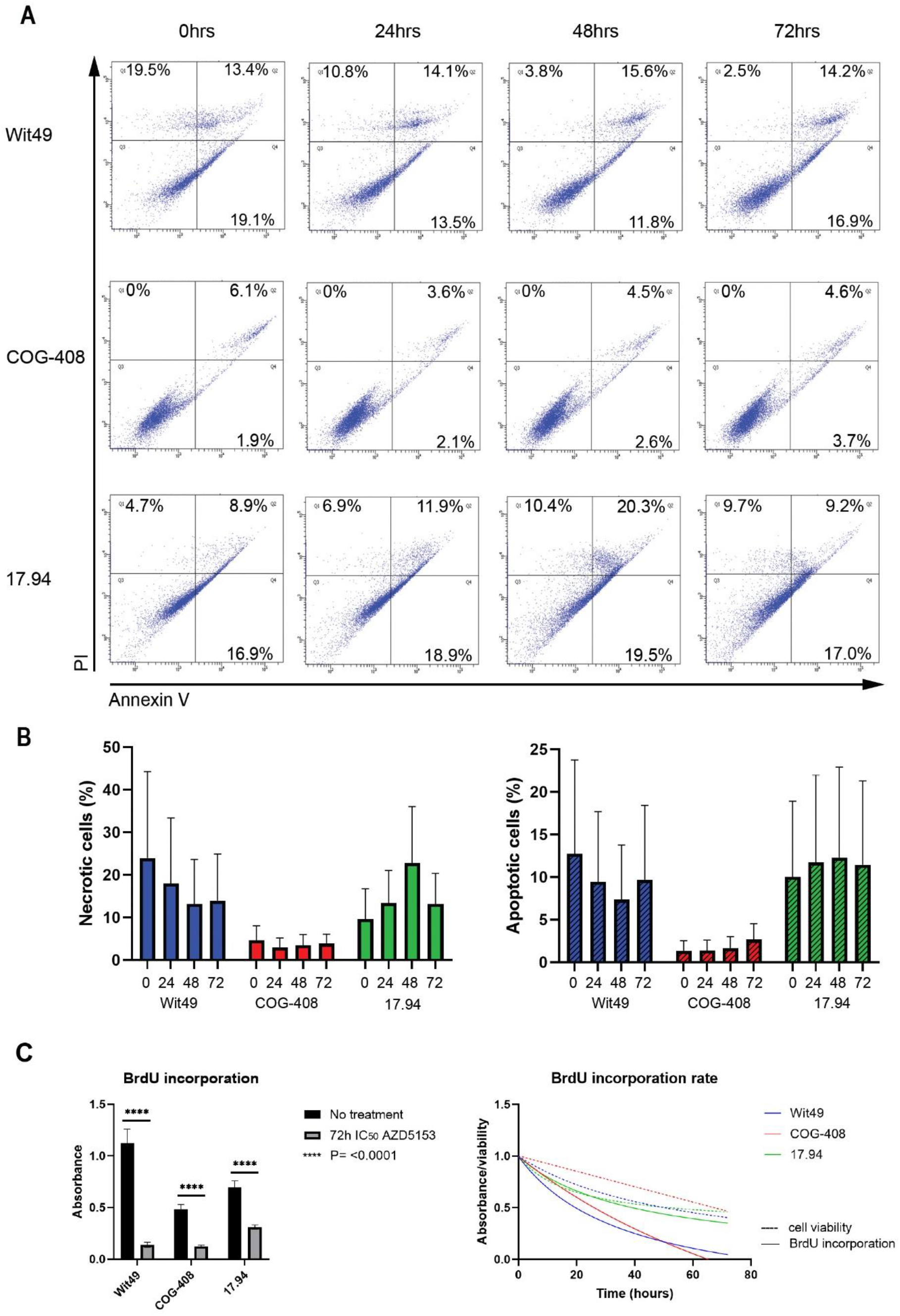
BRD4 inhibition results in apoptosis and significantly suppresses proliferation in Wilms’ tumor cell lines. A. Flow cytometry dot plots showing rates of apoptotic and necrotic cells over time in Wilms’ tumor cells exposed to model-specific IC_50_ concentrations of AZD5153. B. Quantification of necrotic and apoptotic cells over time with model-specific IC_50_ concentrations of AZD5153. C. (Left) BrdU incorporation (as measured by absorbance) in un-treated Wilms’ tumor cells vs. AZD5153 treated cells. A 2-way ANOVA analysis showed significant reduction of BrdU incorporation (P = <0.0001) in the treatment condition across all cell lines. (Right) Rate of BrdU incorporation over time as compared to cell viability upon exposure to model-specific IC_50_ concentrations of AZD5153.

We next considered AZD5153’s effect on radiosensitivity. In the clinical setting, Wilms’ tumor is known to be radiosensitive, with radiation frequently used as a treatment option for advanced and aggressive cases (Tefft et al., 1976, Dome et al., 2015). We chose two Wilms’ tumor cell lines sensitive to radiation and one resistant cell line. We observed a slight additive effect in the radiation+AZD5153 treatment group in the radiosensitive cell lines Wit49 and COG-W-408 (**Fig EV 3**). No effect was observed in the insensitive cell line 17.94.

### Clinically-representative dosing schedule of BRD4 inhibitor AZD5153 abrogates Wilms’ tumor cell growth

In order to explore cellular effects in the presence of clinically achievable levels of AZD5153, microfluidics experiments with anaplastic Wilms’ tumor cell line Wit49 were performed. We used a similar dosing schedule to the recommended phase II dose (RP2D) of 15 mg/kg BID, as determined by clinical trial NCT03205176. For AZD5153, the maximum plasma concentration (Cmax) and concentration at steady state (Css) were estimated to be 110 nM and 58 nM, respectively, with a total AUC of 800 nM*h, when dosed 2 times per day. This dosing schedule was replicated (**Fig 5A**) and cell death and impairment of cell growth was observed as compared to DMSO control (**Fig 5C** and **Video EV 1-2**). Cell confluency (surface area) was measured as a proxy for viable cells and compared between the two groups (**Fig 5B**). At the end of the experiment, we observed 5-fold fewer viable cells in the AZD5153 treatment group. Notably, while some degree of apoptosis and cell death was observed within the treatment condition, suppression of proliferation and anti-adhesion activities were also noted (**Fig 5D**) and appear to be major drivers of the resultant lack of cellular growth as compared to control.

**Figure 5.**
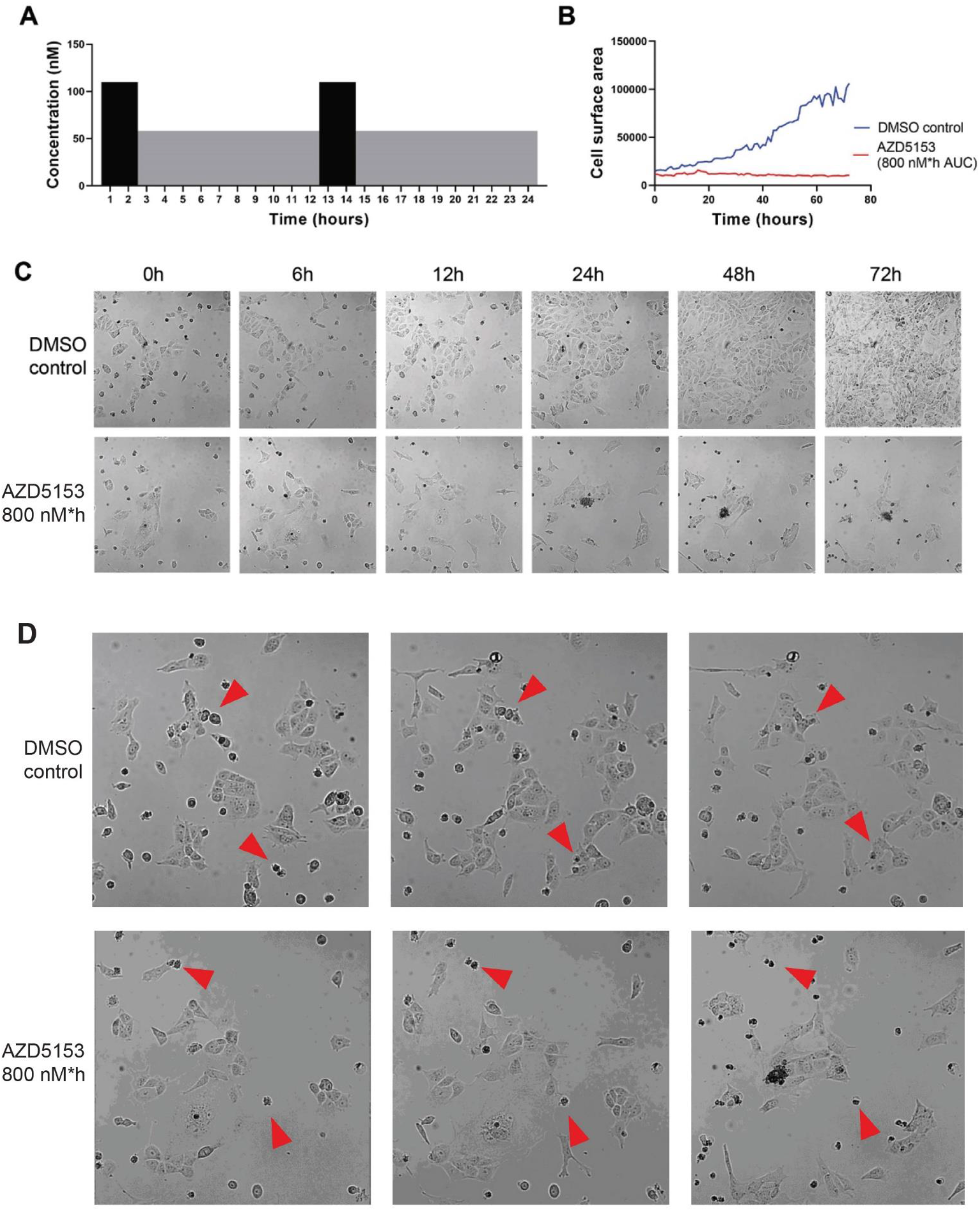
Microfluidics images with clinically representative AZD5153 dosing schedule. A. AZD5153 dosing schedule using clinically achievable drug concentrations (Cmax=110nM, Css=58nM, AUC=800 nM*h). B. Quantification of cell surface area following exposure to AZD5153 or control (DMSO). Fewer cells were observed in the treatment group. C. Time-lapsed images of Wit49 cell culture exposed to AZD5153 or control (DMSO). D. Untreated cells adhere to the plate and proliferate. AZD5153 treated cells are unable to adhere and lack proliferation activity.

Although two separate histological subtypes of Wilms’ tumor patients are seen in the clinic, each with distinct outcomes and response to treatments, only a small handful of differences at the genomic level between the two groups have been observed. The MYCN oncogene is known to be involved in many oncogenic processes and has been associated with the anaplastic subtype of Wilms’ tumor (Williams et al., 2015), which was confirmed within this study. Our data suggests that reducing MYCN levels is a potential therapeutic strategy for management of anaplastic Wilms’ tumor.

Here, we have demonstrated that MYC and MYCN levels in Wilms’ tumor cell lines can be transiently modulated via BET/BRD inhibition with AZD5153, a potent and selective bivalent inhibitor of BRD4, and that this modulation results in decreased cell viability, anti-adhesion activity and suppressed proliferation potential. Additional *in vivo* studies are needed to determine if the observed rebound in MYC levels results in a return to tumor progression in Wilms’ tumor, or if only a brief reduction in MYC protein levels can result in sustained tumor response, as has been seen in other cancer types such as osteogenic sarcoma (Jain et al., 2002). Alternatively, AZD5153 combined with additional targeted therapies may provide synergistic effects against Wilms’ tumors, or might result in an extension to the period of *MYC* and *NMYC* downregulation. This approach should be pursued with caution, however, as sustained oncogene reduction might result in significant toxicities due to the disruption of critical signaling pathways in normal cells.

A limitation to our work is the small number of anaplastic Wilms’ tumor cell lines available for analysis. The research community stands to benefit greatly from the development of additional Wilms’ tumor cell lines, especially those which can be characterized by anaplasia or other high-risk factors. New approaches to Wilms’ tumor cell line development are needed.

AZD5153 has demonstrated a favorable toxicity profile in both mouse and human studies. Our study has demonstrated that AZD5153 treatment can effectively reduce MYC protein levels in Wilms’ tumor models. We conclude that *BRD4* is a relevant target in Wilms’ tumor and that AZD5153 should be further explored as a potential therapeutic for patients with advanced and resistant disease.

## Materials and methods

### Cell lines and patient samples

The COG-W-408 cell line was obtained from the Childhood Cancer Repository (Lubbock, TX) and maintained in Iscove’s Modified Dulbecco’s Medium supplemented with 20% Fetal Bovine Serum, 4mM L-Glutamine and 1X ITS (5 μg/mL insulin, 5 μg/mL transferrin, 5 ng/mL selenium) (all from Gibco, Thermo Fisher Scientific, Waltham, MA). The Wit49 cell line was a gift from Dr. Herman Yeger (Sick Kids Hospital, Toronto, Canada) and was maintained in Dulbecco’s Modified Eagle Medium: Nutrient Mixture F-12 supplemented with 10% Fetal Bovine Serum and 1% Penicillin-Streptomycin solution (all from Gibco, Thermo Fisher Scientific). The 17.94 cell line was purchased from DSMZ bioresource center (Brauchemburg, Germany) and was maintained in Dulbecco’s Modified Eagle Medium supplemented with 20% Fetal Bovine Serum and 1% Penicillin-Streptomycin solution (all from Gibco, Thermo Fisher Scientific). The IM-WT-1, IM-WT-6, IM-WT-10 cell lines and Wilms-8 primary culture were provided by co-author Dr. Brigitte Royer-Pokora (University Hospital Düsseldorf Institute of Human Genetics, Düsseldorf, Germany) and were maintained in Mesenchymal Stem Cell Growth Medium (PT-3001, Lonza, Basel, Switzerland). The HEK293 cell line was purchased from the American Type Culture Collection (Manasses, VA) and maintained in RPMI-1640 cell culture medium supplemented with 10% Fetal Bovine Serum and 1% Penicillin-Streptomycin solution (all from Gibco, Thermo Fisher Scientific). Primary cell culture CF-00108 was a gift from Dr. Renata Veselska (Masaryk University, Brno, Czech Republic) and was maintained in Dulbecco’s Modified Eagle Medium supplemented with 20% Fetal Bovine Serum and 1% Penicillin-Streptomycin solution (all from Gibco, Thermo Fisher Scientific). Primary cell cultures PCB-00007, CF-00136, and CF-00333 were collected from patients enrolled in the CuReFAST tumor banking study at the Children’s Cancer Therapy Development Institute who had given informed consent. Tumor tissue was minced by hand and then processed in a GentleMacs dissociator (130-093-235, Miltenyi Biotec GmbH, Bergisch Gladbach, Germany) according to manufacturer’s protocol. Resultant cultures were maintained in RPMI-1640 cell culture medium supplemented with 10% Fetal Bovine Serum and 1% Penicillin-Streptomycin solution (all from Gibco, Thermo Fisher Scientific). All cell cultures were maintained in a humidified incubator at 37°C supplemented with 5% CO_2_. Short Tandem Repeat validation and mycoplasma testing was performed on all cell cultures (**Table EV1 and Table EV 2**). KT-13 and KT-18 PDX explants were generously provided by Dr. Peter Houghton (University of Texas Health Science Center, San Antonio, TX, USA). A table of Wilms’ tumor models used in this study is found in **Table EV 1**.

### Chemical screens

Discovery screens were conducted using an investigator-selected 60-agent drug screen. Endpoints were per drug IC50 values. Each agent was tested in triplicate at 4 dosage points. Initial drug stocks were diluted in appropriate solvents, then plated onto Nunc™ 384-Well Polystyrene White Microplates (164610, Thermo Fisher Scientific) with DMSO and media-only controls. Drug plating was performed with EpMotion 5075 (Eppendorf, Hamburg, Germany) and Sciclone G3 (PerkinElmer) liquid handling robots. On day 0, cells were trypsinized and added to the drug plates (2000 cells/well) with a Multi-Flo liquid dispenser (BioTek, Winooski, VT), then placed in a humidified incubator for 72 hours at 37°C supplemented with 5% CO2. Cell viability (ATP) was measured using the CellTiter-Glo^®^ Luminescent Cell Viability Assay (G7573, Promega, Madison, WI) following manufacturer’s protocol. Output was read with a Synergy HT plate reader (BioTek).

Follow on drug screens were conducted with a Tecan D300e digital dispenser (Hewlett Packard, Palo Alto, CA). BET/BRD inhibitors JQ1, I-BET-762, BI5352, LY20094, and AZD5153 were purchased from Selleck Chemicals (Houston, TX). Initial drug stocks were mixed with DMSO to a 10 mM concentration. Cells were plated (2000 cells/well) onto Nunc™ 384-Well Polystyrene White Microplates (164610, Thermo Fisher Scientific) in culture-appropriate media and allowed to adhere for 12 hours. For IC50 experiments, drugs were applied at 10 different concentrations (4.3 nM to 10 μM, logarithmic scale) in triplicate and then placed in a humidified incubator for 72 hours at 37°C supplemented with 5% CO2. Cell viability (ATP) was measured using the CellTiter-Glo^®^ Luminescent Cell Viability Assay (G7573, Promega) as described above. Values were normalized to background (media only) and control (untreated cells and media). Nonlinear regression curves and IC50 values were determined using Graphpad Prism 8.2.1 software.

### Genetic sequencing

Wilms’ tumor samples were sequenced at the Beijing Genomics Institute (Cambridge, MA) with 100x coverage paired-end whole exome for DNA, and 40M paired-end deep transcriptome for RNA. Samples were tested using an Agilent 2000 bioanalyzer (Agilent Technologies, Santa Clara, CA). Library was constructed using the BGISeq-500 library construction protocol. At least 12G of data per sample was generated for analysis. Whole exome sequencing data was analyzed for the presence of somatic point mutations, somatic functional and structural mutations, potential germline mutations, polynucleotide insertions and deletions, and gene copy number variation. Somatic mutations, variations, and indels were called using Genome Analysis Toolkit (GATK) Version 4.0 with strict calling criterion (Tumor logarithm of odd (TLOD) scores > 6.3). Gene copy number variations were identified using Samtools and VarScan2 quantified as a log ratio of tumor copy to normal copy using the GRCh38 human reference genome. RNA sequencing data was analyzed for gene expression and gene fusion events. Transcriptome data was aligned to STAR-derived human transcriptome from GRCh38 human reference genome. Normalized gene expression was quantified using RSEM. Region-specific (kidney) gene expression data was accessed from the Genotype–Tissue Expression (GTEx) project to serve as a population normal and to identify underexpressed and overexpressed genes. The data discussed in this publication have been deposited in NCBI’s Gene Expression Omnibus (Edgar et al., 2002) and are accessible through GEO Series accession number GSE156065 (https://www.ncbi.nlm.nih.gov/geo/query/acc.cgi?acc=GSE156065).

### TARGET sequencing data

The results published here are in part based upon data generated by the Therapeutically Applicable Research to Generate Effective Treatments (https://ocg.cancer.gov/programs/target) initiative, phs000218. The data used for this analysis are available at https://portal.gdc.cancer.gov/projects. Wilms’ tumor data can be found under the study accession number phs000471. Specimen clinical information was gathered from the TARGET Data Matrix (https://ocg.cancer.gov/programs/target/data-matrix). Whole exome and gene expression sequencing data from 125 anaplastic and high risk (relapsed) Wilms’ tumor patients was analyzed. Paired-end whole exome data was collected on the HiSeq 2000 sequencing system (Illumina, San Diego, CA). Gene expression data was collected using the Gene Chip^®^ Human Genome U133 Plus 2.0 Array (Affymetrix, Santa Clara, CA) protocol. FASTQ files were downloaded and analyzed as described above.

### Overall survival analysis

Gene expression transcripts per million (TPM) levels and clinical data were collected from TARGET study samples as described above. The significance of variation in overall survival time with *MYC* TPM and *MYCN* TPM was assessed with a proportional hazards models without and with adjustment for age, sex and histology. The hazard ratios, standard errors, 95% percent confidence intervals and p-values were displayed. Survival distributions were described with Kaplan-Meier curves. All statistical testing was twosided with a significance level of 5%. Corrections for multiple comparisons were not applied. SAS Version 9.4 was used throughout.

### Dendrogram and cluster analysis

Hierarchical agglomerative clustering analysis was performed on a panel of *MYC*-related gene expression levels as measured by transcripts per million (TPM) using Python’s seaborn library. The TPM values were transformed using log2 (x+1) before clustering. Wilms’ tumor samples used within the analysis included anaplastic and non-anaplastic cell lines (n=10), anaplastic PDX explants (n=2) and median data from the NCI TARGET study (n=135). Region-specific (kidney) gene expression data from Genotype-Tissue Expression (GTEx) project and median values from the samples were also plotted in the clustering analysis. The Euclidean distance metric was used for clustering. Axes were color coded to highlight additional information about the data such as sample histology status and *MYC*-related gene function for genes of interest.

### Western blot analysis

Cells were washed with cold phosphate buffered saline and mechanically extracted in 1x radioimmunoprecipitation assay (RIPA) lysis and extraction buffer supplemented with complete protease inhibitor and phosphatase inhibitor cocktail (89901 and 78441, Thermo Fisher Scientific). After incubation at 4°C for 30 min, cells were centrifuged (12,000g for 5 min at 4°C), and supernatant was collected. Protein was quantified using the Pierce BCA assay kit (23224, Thermo Fisher Scientific). Fifty micrograms of protein from each sample was loaded and separated in 7.5% SDS-PAGE gel (4561024, Bio-Rad, Hercules, CA) and transferred onto a 0.2 mm PVDF membrane using the wet transfer method (90 V for 90 min). Primary C-MYC tag antibody (PA1-981, Thermo Fisher Scientific) and N-*MYC* Polyclonal antibody (PA5-17403, Thermo Fisher Scientific) were diluted 1:500 in 5% non-fat powdered milk and 1X TBST, then placed on a rocker overnight at 4°C. After overnight incubation, PVDF membrane was washed 3 times in 1X Tris-buffered saline + Tween20 (TBST) (15 minutes each on medium speed rocker at room temperature). Peroxidase labeled anti-rabbit secondary antibody (PI-1000, Vector laboratories, Burlingame, CA) was diluted 1:5000 in 5% non-fat powdered milk and 1X TBST, then placed on a rocker at room temperature for 1 hour. After secondary antibody incubation, the PVDF membrane was washed as before. Proteins were detected with Clarity Western ECL Substrate (1705061, Bio-Rad) according to manufacturer’s protocol and read on an IVIS Lumina Imaging System (PerkinElmer, Waltham, MA), with an exposure time of 5-10 seconds. Molecular weights were determined using Precision Plus Protein™ Dual Color Standards protein ladder (1610394, Bio-Rad). All experiments were conducted in triplicate. Protein levels were quantified using Photoshop 2021 software and analyzed with Graphpad Prism 8.2.1 software.

### Annexin V staining experiments

Each of 3 Wilms’ tumor cell lines were plated into 4, 10cm tissue culture plates (2 million cells each). Model-specific IC50 doseages of AZD5153 were added over a 72-hour time period. Culture plates were divided into 4 treatment conditions: 0 hours (no treatment), 24 hours, 48 hours and 72 hours of drug exposure. An additional “no treatment” plate was used as an unstained control. Cells were stained with the Annexin V-FITC Apoptosis Staining/Detection Kit (ab14085, Abcam, Cambridge, MA) and visualized by flow cytometry on a BD FACSAria II (Becton Dickinson Biosciences, Franklin Lakes, NJ). Briefly, 5 x 10^5^ cells were collected by trypsinization, then re-suspended in ANXA5 binding buffer. Annexin V-FITC and Propidium Iodide stains were incubated with cells for 5 minutes at room temperature in the dark before performing flow cytometry by fluorescent-activated cell sorting (FACS) on a BD FACSAria. Analysis of the data was done with BD FACSAria and Graphpad Prism 8.2.1 software.

### BrdU incorporation experiments

Three Wilms’ tumor cell lines were exposed to model-specific IC50 concentrations of AZD5153 for 0, 24, 48 and 72 hours. Cell viability was measured using the using the CellTiter-Glo^®^ Luminescent Cell Viability Assay (G7573, Promega) following manufacturer’s protocol. BrdU incorporation was measured using the BioVision BrdU proliferation plate reader assay kit (306-200, BioVision, Milpitas, CA) following manufacturers protocol. Viability (luminescense) and BrdU incorporation (absorbance) was visualized on a Synergy HT plate reader (BioTek). Data was analyzed using Graphpad Prism 8.2.1 software.

### Microfluidics experiments

Microfluidics experiments were conducted with the CellASIC^®^ ONIX2 microfluidic system (EMDMillipore, Burlington, MA) following manufacturer’s protocols. Briefly, 10,000 cells were loaded into each chamber of a CellASIC ONIX M04 switching plate and allowed to incubate overnight in complete media. The next day, stock solutions of 110nM AZD5153 and 58nM AZD5153, DMSO controls and additional complete media were loaded into the microfluidic plate, which was then placed inside a XLMulti S1 incubator (Pecon, Erbach, Germany) mounted onto a LSM800 confocal inverted laser scanning microscope (Zeiss, Oberkochen, Germany). Cells were kept at 37°C and supplemented with 5% CO2. Images were captured every 30 minutes over a 72-hour period with Zen2.5 software. Drug dosing schedule replicated human AUC concentrations as published (AstraZeneca data). Cell surface area was quantified using FIJI ImageJ software.

### *In vitro* radiation experiments

All experiments were conducted in triplicate and results averaged. Two thousand cells/well were seeded in complete media onto Nunc™ 96-well white-walled tissue culture plates (136101, Thermo Fisher Scientific), using a Multi-Flo (BioTek) liquid dispenser. Plates were placed in the center of the top shelf of a MX-20 X-ray cabinet (Faxitron, Tucson, AZ) and radiated for 30 minutes every 12 hours over a 72-hour period at 35 kV for a total delivered X-ray dose of 61.5 Gy. A 37° deltaphase isothermal warming pad was placed directly under the plate to prevent cells from cooling. For the drug treatment conditions, cells were plated in the presence of IC50 concentrations of AZD5153. At 72 hours, cell viability was measured using the CellTiter-Glo^®^ Luminescent Cell Viability Assay (G7573, Promega) according to manufacturer’s protocol. Luminescence was read on a Synergy HT plate reader (BioTek). Output data was normalized against an untreated plate for control (cells and media) and background (media). P-values for radiation experiments were determined by 2-tailed Welch’s t-test.

## Data availability

The datasets produced in this study are available in the following databases:

- RNA-Seq data: Gene Expression Omnibus, accession number GSE156065 (https://www.ncbi.nlm.nih.gov/geo/query/acc.cgi?acc=GSE156065).
- TARGET RNA-seq data: Therapeutically Applicable Research to Generate Effective Treatments (https://ocg.cancer.gov/programs/target) initiative, phs000218. Wilms’ tumor data can be found under the study accession number phs000471. (https://portal.gdc.cancer.gov/projects). Clinical data is stored at (https://ocg.cancer.gov/programs/target/data-matrix).

## Compliance with ethical standards

All human tissue samples for primary cell generation were reviewed and approved by the Children’s Cancer Therapy Development Institute’s Institutional Review Board (Advarra, protocol # cc-TDI-IRB-1) and collected through the Cancer Registry for Familial and Sporadic Tumors (CuReFAST) tumor banking study. All patients enrolled in CuReFAST provided informed consent. Patient data and clinical and pathologic information are maintained in a de-identified, encrypted and secure database.

## Acknowledgements

This work was supported by research grants from the Rally and CURE Childhood Cancer Foundations, The Truth 365, Infinite Love for Kids Fighting Cancer, Joey’s Wings, Alex’s Army, Unravel Pediatric Cancer, The Bozeman 3, Hope from Harper and the Wilms’ tumor patient/family community. This study is dedicated to Stellablue, anaplastic Wilms’ tumor survivor. We are grateful to Dr. Herman Yeger (Sick Kids Hospital, Toronto, CA), Dr. Renata Veselska (Masaryk University, Brno, Czechia) and Dr. Peter Houghton (Greehey Children’s Cancer Research Institute, TX, USA) for providing samples. Thank you to Dr. Naomi Pode-Shakkad and Dr. Michael Ortiz for inspiration, support and helpful discussion.

## Author contributions

ADW was involved in the design of this study and the experiments, performed experiments, and wrote the manuscript. NEB and CK designed experiments, analyzed data and participated in writing the manuscript. JM performed the survival analysis. KF provided statistical review and analysis. RP helped with experiments and figure development. ML, KM, GS and NB helped with genetic analysis. BRP and RV provided resources and critical discussion. All authors read and approved the final manuscript.

## Expanded View Figure Legends

**Expanded View Figure 1.**
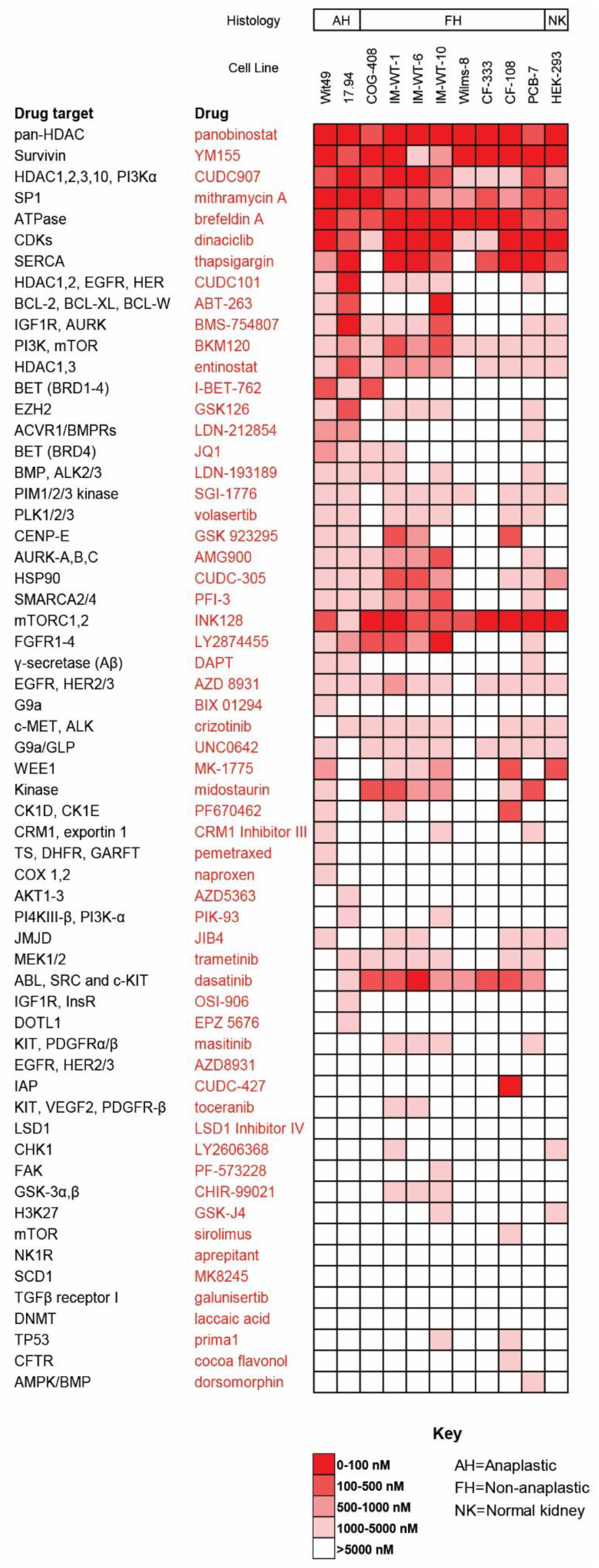
V4 drug screen. Expanded (60-compound) drug screen results against 2 anaplastic Wilms’ tumors, 8 non-anaplastic Wilms’ tumors, and 1 normal kidney cell line.

**Expanded View Figure 2.**
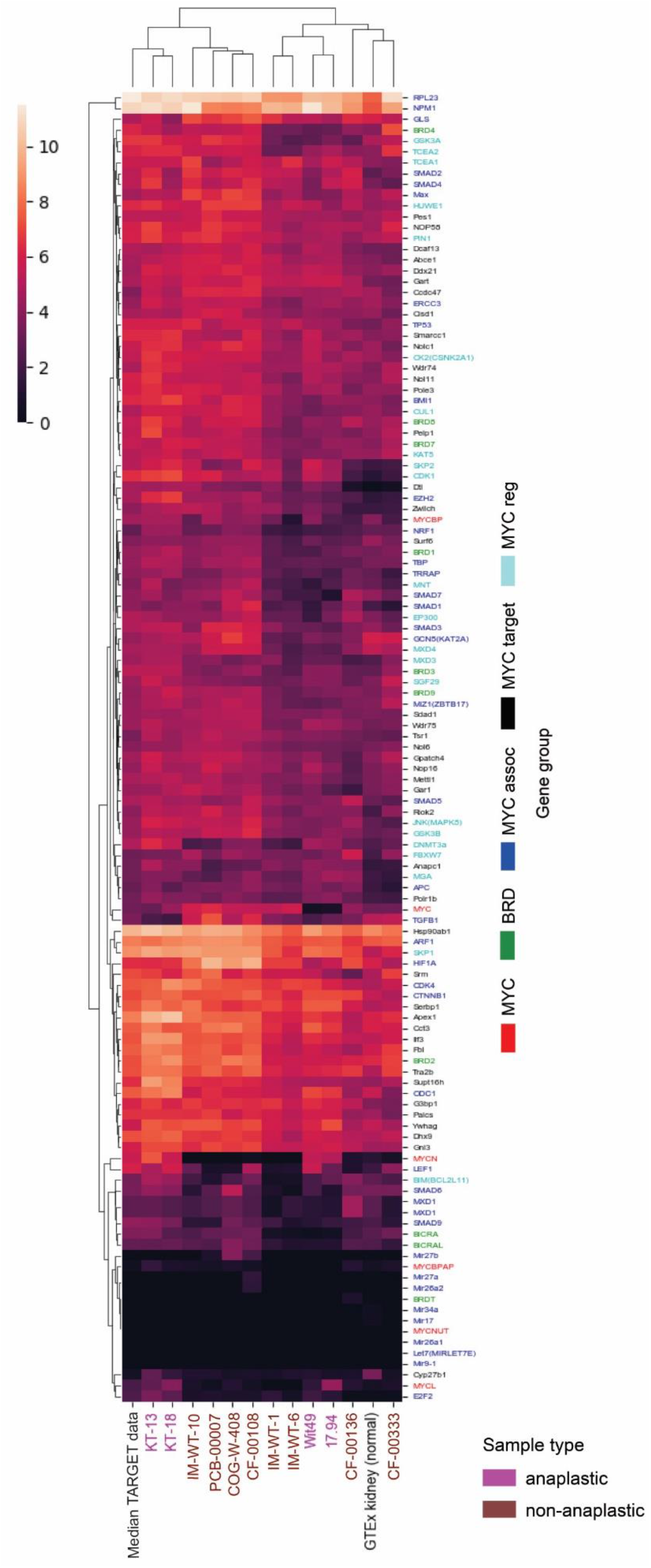
Dendrogram heatmap of MYC genes in Wilms’ tumors. Cluster analysis and dendrogram heatmap of Wilms’ tumor gene expression levels (TPM) across an expanded panel of MYC related genes.

**Expanded View Figure 3.**
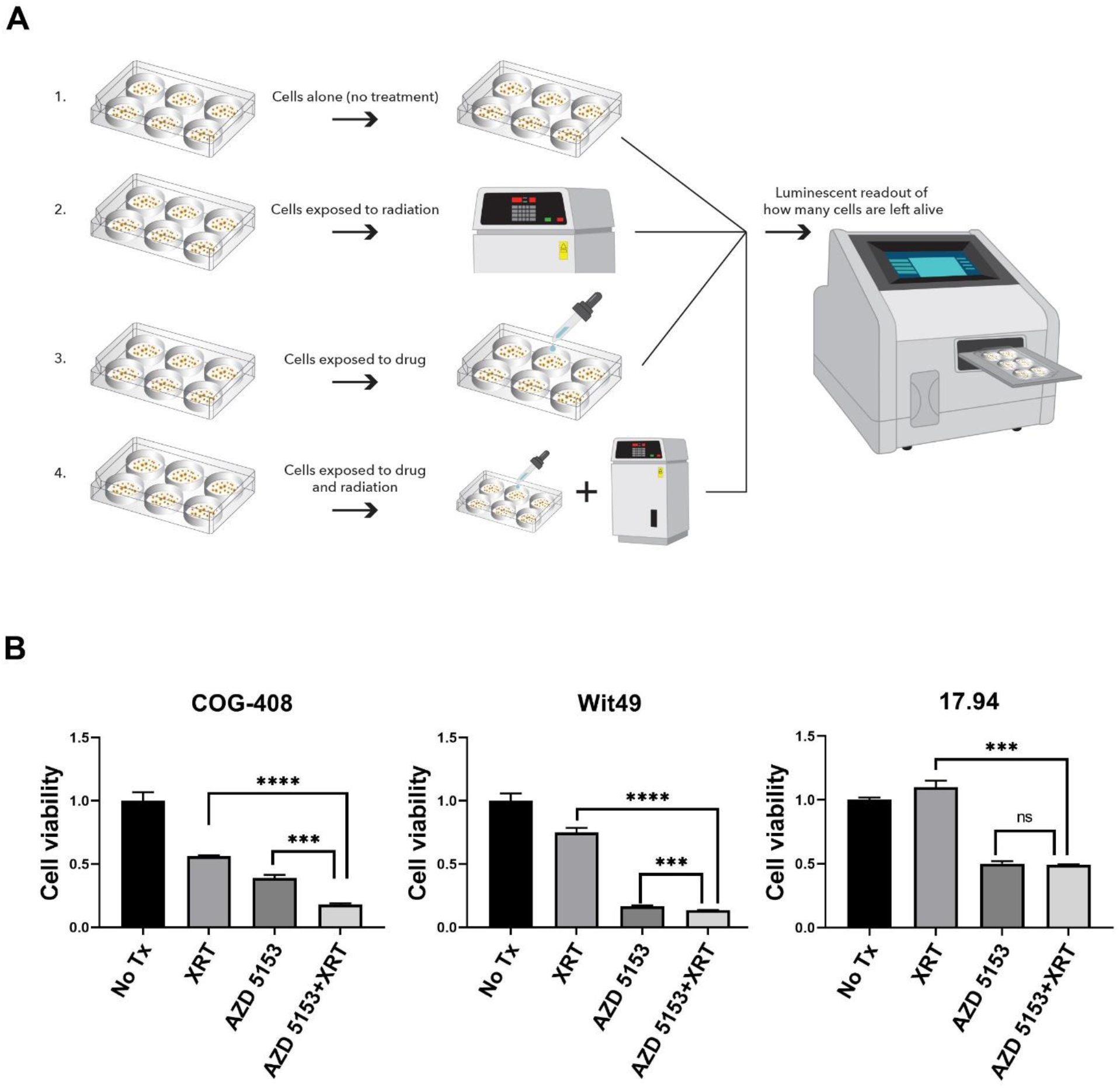
Radiation+AZD5153 combination against Wilms’ tumor cell lines. A. Schematic representation of experimental procedure. Wilms’ tumor cell lines COG-W-408, Wit49 and 17.94 were treated with 4 conditions: no treatment, radiation only, drug only, and radiation+drug. Cell viability was measured at 72 hours. B. Combination of radiation (X-ray) and drug exposure results in a significant decrease of Wilms’ tumor cell viability in radiosensitive cell lines Wit49 and COG-W-408. Non-radiosensitive cell line 17.94 responds to AZD5153 treatment but does not show a significantly increased response to the radiation/drug combination.

## Expanded View Table Legends

**Expanded View Table 1.** Wilms’ tumor sample information. Clinical data and validation results of cell lines, PDX models and primary cultures used within this study: 4 anaplastic Wilms’ tumors, 9 non-anaplastic Wilms’ tumors, and 1 normal kidney model.

| Name | Model Type | Histology | Sex | Age (years) | STR result | Mycoplasma test |
| --- | --- | --- | --- | --- | --- | --- |
| Wit49 | Cell line | Anaplastic | Female | 2 | No reference profile, No match | Negative |
| 17.94 | Cell line | Anaplastic | Female | 4 | Matched reference profile | Negative |
| KT-13 | PDX explant | Anaplastic | Female | 3 | Not tested | Not tested |
| KT-18 | PDX explant | Anaplastic | Male | 5 | Not tested | Not tested |
| COG-W-408 | Cell line | Non-anaplastic | Unknown | 9 | No reference profile, No match | Negative |
| IM-WT-1 | Cell line | Non-anaplastic | Female | 1 | No reference profile, No match | Negative |
| IM-WT-6 | Cell line | Non-anaplastic | Male | 1 | No reference profile, No match | Negative |
| IM-WT-10 | Cell line | Non-anaplastic | Female | 2 | No reference profile, No match | Negative |
| Wilms-8 | Primary cell culture | Non-anaplastic | Male | 1 | No reference profile, No match | Negative |
| CF-00108 | Primary cell culture | Non-anaplastic | Female | 1 | No reference profile, No match | Negative |
| CF-00136 | Primary cell culture | Non-anaplastic | Male | 24 | No reference profile, No match | Negative |
| CF-00333 | Primary cell culture | Non-anaplastic | Male | 38 | No reference profile, No match | Negative |
| PCB-00007 | Primary cell culture | Non-anaplastic | Female | 5 | No reference profile, No match | Negative |
| HEK293 | Cell line | Normal kidney | Female | fetus | Matched reference profile | Negative |

**Expanded View Table 2.** Short Tandem Repeat (STR) profiles of Wilms’ tumor samples. STR profiles for study models used within this study.

| Name | D3S1358 | TH01 | D21S11 | D18S51 | Penta_E | D5S818 | D13S317 | D7S820 | D16S539 | CSF1PO | Penta_D | AMEL | vWA | D8S1179 | TPOX | FGA |
| --- | --- | --- | --- | --- | --- | --- | --- | --- | --- | --- | --- | --- | --- | --- | --- | --- |
| Wit49 | 17, 17 | 9.3, 9.3 | 29, 32.2 | 15, 15 | 13, 13 | 12, 13 | 11, 12 | 8, 11 | 12, 12 | 11, 12 | 11, 12 | X, X | 16, 18 | 10, 10 | 8, 8 | 25, 25 |
| 17.94 | 16, 16 | 8, 9.3 | 29, 29 | 14, 17 | 7, 11 | 11, 11 | 8, 8 | 11, 11 | 11, 11 | 11, 12 | 9, 11 | X, X | 14, 16 | 10, 13 | 8, 8 | 22, 23 |
| COG-408 | 15, 18 | 9.3, 9.3 | 28, 28 | 10, 17 | 12, 15 | 9, 12 | 8, 12 | 11, 12 | 9, 11 | 12, 13 | 9, 13 | X, X | 16, 19 | 11, 12 | 8, 8 | 24, 25 |
| IM-WT-1 | 14, 17 | 9.3, 9.3 | 30, 31 | 15, 18 | 5, 14 | 12, 13 | 11, 13 | 9, 14 | 11, 14 | 10, 10 | 13, 13 | X, X | 14, 19 | 12, 14 | 8, 9 | 22, 25 |
| IM-WT-6 | 14, 17 | 9.3, 9.3 | 30, 32 | 15, 18 | 5, 14 | 12, 13 | 11, 13 | 9, 14 | 11, 14 | 10, 10 | 13, 13 | X, X | 14, 19 | 12, 14 | 8, 9 | 22, 25 |
| IM-WT-10 | 17, 17 | 8, 8 | 29, 30 | 14, 16 | 7, 7 | 10, 12 | 12, 12 | 11, 12 | 9, 10 | 11, 11 | 10, 13 | X, Y | 15, 18 | 10, 15 | 8, 11 | 22, 24 |
| Wilms-8 | 16, 18 | 8, 8 | 29, 33.2 | 12, 12 | 12, 17 | 12, 13 | 8, 9 | 8, 10 | 13, 13 |  | 10, 12 | X, Y | 18, 18 | 8, 13 | 8, 9 | 20, 21 |
| CF-00108 | 14, 15 | 6, 9 | 31, 31.2 | 14, 18 | 7, 10 | 11, 11 | 8, 10 | 9, 9 | 11, 13 | 9, 14 | 12, 12 | X, X | 17, 18 | 12, 14 | 8, 9 | 21, 22 |
| CF-00136 | 16, 17 | 6, 9.3 | 28, 30 | 13, 21 | 11, 12 | 7, 13 | 9, 13 | 10, 10 | 10, 11 | 12, 12 | 11, 12 | X, Y | 17, 18 | 13, 13 | 8, 8 | 20, 22 |
| CF-00333 | 14, 15 | 6, 9.3 | 29, 29 | 14, 14 | 7, 12 | 9, 12 | 9, 11 | 9, 11 | 11, 12 | 10, 10 | 11, 12 | X, Y | 16, 20 | 13, 15 | 8, 11 | 20, 21 |
| PCB00007 | 16, 18 | 7, 9.3 | 29, 32.2 | 17, 18 | 15, 17 | 11, 13 | 11, 12 | 10, 12 | 9, 11 | 11, 13 | 5, 12 | X, X | 18, 19 | 11, 15 | 9, 11 | 19, 23 |
| HEK293 | 15, 17 | 7, 9.3 | 28, 30.2 | 17, 18 | 7, 15 | 8, 8 | 12, 14 | 11, 12 | 9, 13 | 12, 12 | 9, 10 | X, Y | 16, 19 | 12, 14 | 11, 11 | 23, 23 |

## Expanded View Movie Legends

**Expanded View Movie 1 and 2 - Microfluidics videos showing cell killing and growth inhibition of Wilms’ tumor cells in the presence of a clinically representative AZD5153 dosing schedule.**

A. 72-hour microfluidics experiment showing apoptosis and proliferation suppression of Wit49 anaplastic Wilms’ tumor cells in the presence of AZD5153 at clinically representative doses.

B. DMSO control (no drug) condition showing normal Wit49 cell growth.

